# Praziquantel inhibits *Caenorhabditis elegans* development and species-wide differences might be cct-8-dependent

**DOI:** 10.1101/2023.05.17.541211

**Authors:** Janneke Wit, Clayton M. Dilks, Gaotian Zhang, Karen S. Kim Guisbert, Stefan Zdraljevic, Eric Guisbert, Erik C. Andersen

## Abstract

Anthelmintic drugs are used to treat parasitic roundworm and flatworm infections in humans and other animals. *Caenorhabditis elegans* is an established model to investigate anthelmintics used to treat roundworms. In this study, we use *C. elegans* to examine the mode of action and the mechanisms of resistance against the flatworm anthelmintic drug praziquantel (PZQ), used to treat trematode and cestode infections. We found that PZQ inhibited development and that this developmental delay varies by genetic background. Interestingly, both enantiomers of PZQ are equally effective against *C. elegans*, but only the left-handed PZQ (S-PZQ) is effective against schistosome infections. We conducted a genome-wide association mapping with 74 wild *C. elegans* strains to identify a region on chromosome IV that is correlated with differential PZQ susceptibility. Five candidate genes in this region: *cct-8, znf-782, Y104H12D*.*4, Y104H12D*.*2*, and *cox-18*, might underlie this variation. The gene *cct-8*, a subunit of the protein folding complex TRiC, has variation that causes a putative protein coding change (G226V), which is correlated with reduced developmental delay. Gene expression analysis suggests that this variant correlates with slightly increased expression of both *cct-8* and *hsp-70*. Acute exposure to PZQ caused increased expression of *hsp-70*, indicating that altered TRiC function might be involved in PZQ responses. To test if this variant affects development upon exposure to PZQ, we used CRISPR-Cas9 genome editing to introduce the V226 allele into the N2 genetic background (G226) and the G226 allele into the JU775 genetic background (V226). These experiments revealed that this variant was not sufficient to explain the effects of PZQ on development. Nevertheless, this study shows that *C. elegans* can be used to study responses to PZQ to identify mode of action and resistance mechanisms. Additionally, we show that the TRiC complex requires further evaluation for PZQ responses in *C. elegans*.

## 1. INTRODUCTION

Praziquantel (PZQ) is an anthelmintic drug used to treat platyhelminth infections, including schistosomiasis in humans [1,2]. Schistosomiasis is caused by an infection with one of six schistosome species (*Schistosoma guineensis, S. haematobium, S. intercalatum, S. japonicum, S. mansoni*, and *S. mekongi*) and affected over 250 million people in 2021 alone [3]. Preventative treatment with PZQ in areas where schistosomes are endemic is part of the World Health Organisation’s recommendation for control and ultimate elimination of human schistosomiasis [3]. PZQ is the only drug currently available to treat infections in humans, and PZQ resistance in schistosomes is not widely reported at present [4,5]. However, with the increase in mass drug administration (MDA) programs, selection for PZQ resistance increases and will cause decreased efficacy over time. Indeed, a study of genetic diversity in *S. mansoni* after MDA showed that long-term PZQ exposure selected for putative resistance loci across the genome, although these signatures of selection did not appear linked to a resistance phenotype [5].

Because of the threat that resistance poses for sustained treatment efficacy, it is important to study the genetic basis of resistance, both to identify resistant populations and to prevent their spread [6]. Calcium ion transporters have long been the focus of PZQ mode of action and resistance studies [7–10], and recently, the transient receptor potential melastatin ion channel in *S. mansoni* (*Sm*.TRPM_PZQ_) has been identified as a molecular target of PZQ and linked to variation in PZQ sensitivity [11,12]. However, this identification hinged on one laboratory selected line, and the involvement of *Sm*.TRPM_PZQ_ in resistance against PZQ treatment has not yet been corroborated in natural populations. Importantly, multiple mechanisms can lead to resistance, and more studies are needed to understand the genetic basis of resistance in natural populations [5].

Conducting studies on natural genetic variation and its role in helminth responses is complicated because of their host-dependent life cycles and a limited experimental toolbox [13]. The free-living nematode *Caenorhabditis elegans* is a well established model to study anthelmintic modes of action and resistance [14–19] because of its tractable life cycle, small and well characterized genome, and the availability of genome-editing tools [20]. Although *C. elegans* is a nematode and PZQ is typically used to target platyhelminth infections or external parasites, *C. elegans* has previously been used to test hypothesis regarding PZQ action, including heterologous expression of voltage-operated Ca^2+^ channels [21].

In this study, we explored whether *C. elegans* can be used as a model to study the natural variation of susceptibility to praziquantel. We performed dose-response analyses with the racemic mixture as well as the right- and left-handed enantiomers, given that only the left-handed enantiomer ((*S*)-PZQ) is effective at fighting schistosome infections *in vivo* and *in vitro* [reviewed in 22]. Additionally, we conducted a genome-wide association mapping with 74 wild *C. elegans* strains to detect regions of the genome that are correlated with variation in PZQ susceptibility. We used CRISPR-Cas9 genome editing as well as gene expression analysis to study the role of the candidate gene *cct-8* in PZQ susceptibility. Our results show that *C. elegans* development is affected by PZQ. In the future, the identification of candidate genes in *C. elegans* and their orthologs in schistosomes could aid the study of PZQ resistance of natural schistosome populations.

## 2. MATERIALS AND METHODS

### 2.1 Strains

Animals were grown on nematode growth media agarose (NGMA) plates with 1% agar and 0.7% agarose and were fed *E. coli* OP50 bacteria prior to drug-response testing [23]. The laboratory strain N2 and wild strains (CB4856, DL238, and JU775) from the *C. elegans* Natural Diversity Resource (CeNDR) were used to study the responses to multiple doses of praziquantel (PZQ). A total of 74 *C. elegans* strains were used for genome-wide association mapping, including CB4856, DL238, JU775, and N2. Two allele-replacement strains were generated and tested: ECA485 *cct-8*(*ean8*) in the N2 background and ECA601 *cct-8*(*ean39*) in the JU775 background.

### 2.2 High-throughput PZQ response assay

The phenotypic response to PZQ was measured as described previously [15,24–26]. In summary, the strains were cultured for four generations in uncrowded conditions to avoid the induction of dauer, bleach synchronized, and titered in K medium [27] at a concentration of one embryo per µL for a total volume of 50 µL per well of a 96-well plate. The day after bleach synchronization, hatched L1s were fed *E. coli* HB101 bacterial lysate (Pennsylvania State University Shared Fermentation Facility, State College, PA; [28]). After feeding, nematodes were grown at 20°C for 48 hours with constant shaking. Three L4 larvae were then sorted into new microtiter plates containing K medium, 10 mg/mL HB101 lysate, 50 μM kanamycin, and either 1% DMSO or PZQ dissolved in 1% DMSO. For the four strain assay, 1 mM PZQ racemate was dissolved in 1% DMSO. For the two strain assay, PZQ racemate, (*S*)-PZQ, and (*R*)-PZQ concentrations were 0.25, 0.5, 1, 2, and 3 mM in 1% DMSO. After sorting, animals were cultured and allowed to reproduce for 96 hours at 20ºC with constant shaking. For accurate nematode length measurements, the samples were treated with 50 mM sodium azide (in M9) to straighten their bodies before analysis using the COPAS BIOSORT. The COPAS BIOSORT is a large particle flow measurement device, which measures time of flight (TOF) and extinction (EXT) of objects passing through the flow cell using laser beams. TOF, representing animal length, and EXT, representing optical density of the animal, are both proxies for animal development because animals get longer and more dense during development. For this study, animal optical density is corrected for animal length for each object in each well, which provides a normalized developmental phenotype (norm.EXT). If praziquantel, either the racemate or the enantiomers, negatively affects animals, they are expected to have a smaller norm.EXT. Raw phenotypic data were processed for outliers and analyzed using the R package *easysorter* [29] as described previously [15]. The reported phenotype is median norm.EXT values for each replicate well. For dose responses, all phenotypic values were normalized by deducting the average median norm.EXT value in PZQ conditions from the average median norm.EXT value in control (1% DMSO) conditions. To test if the effects of PZQ and its enantiomers on development differed, a two-way ANOVA and *post hoc* Tukey HSD test were used, including the interaction between drug type and strains (R package *rstatix*, version 0.7.0).

### 2.3 Genome-wide association mapping

Normalized median optical density measurements were collected from populations of 74 wild *C. elegans* strains in control (1% DMSO) or praziquantel (1 mM PZQ in 1% DMSO) conditions using the high-throughput PZQ response assay described above. A genome-wide association mapping was performed using the phenotypes of the 74 wild strains by running the *NemaScan* pipeline (https://github.com/AndersenLab/NemaScan). Single-nucleotide variants (SNVs) that were present in fewer than 5% of the tested strains were removed from the analysis. SNVs with calculated *p*-values greater than the Bonferroni-corrected significance threshold (alpha = 0.05) were considered significant.

### 2.4 eQTL - mediation

To test if expression of genes in the quantitative trait region (QTL) detected with genome-wide association mapping affected the responses to PZQ across 74 wild *C. elegans* strains, we conducted mediation analysis [as described in 30]. In short, we used the genotype at the QTL peak for normalized median optical density measurements, transcript expression traits that have eQTL overlapped with the phenotype QTL, and the phenotypes response as input to perform mediation analysis using the *medTest()* function and 1000 permutations for p-value correction in the R package *MultiMed* (v2.6.0) (https://bioconductor.org/packages/release/bioc/html/MultiMed.html).

### 2.5 Expression of *hsp* genes and *cct-8* after 96 hours of development in PZQ or control conditions

To quantify expression of *cct-8* and heat shock proteins *hsp-16*.*2* and *hsp-70*, the HTA assay described above was adjusted to generate sufficient numbers of nematodes for RNA extraction. Instead of 96-well plates, 12-well plates (Genesee, 25-101) were used. After four generations of plate culturing in uncrowded conditions and bleach synchronization, embryos were titered in K medium at a concentration of half to one embryo per µL for a total volume of 1 mL per well. The day after bleach synchronization, hatched L1s were fed HB101 bacterial lysate. After feeding, nematodes were grown at 20°C for 48 hours with constant shaking. Sixty L4 larvae were then transferred into new 12-well plates containing K medium, 10 mg/mL HB101 lysate, 50 μM kanamycin, and either 1% DMSO or 1 mM PZQ dissolved in 1% DMSO. Animals were then cultured and allowed to reproduce for 96 hours at 20ºC with constant shaking. After 96 hours, the content of the wells was collected, washed with M9 twice, before 1 mL of Trizol (Thermo Fisher, 15596018) was added to the samples. The samples were then split into two technical replicates each and Trizol added to 1 mL total per microfuge tube. The samples were frozen at −80ºC until RNA extraction.

RNA was extracted for one of the technical replicates per biological sample as follows. To disrupt the tissue, 100 µL sand (Sigma, 274739) was added and the samples vortexed vigorously for ten minutes at room temperature (RT). 200 µL chloroform was then added and the sample vortexed for three minutes and spun at full speed for three minutes. The aqueous layer (∼500 µL) was transferred to a new tube and 500 µL of isopropanol was added. Samples were briefly vortexed and incubated for eight minutes at RT before being transferred to ice for two minutes. Samples were then centrifuged at full speed for ten minutes and the supernatant removed. 1 mL freshly made 75% ethanol was added and the samples vigorously vortexed and centrifuged at full speed for three minutes. After removing the supernatant the samples were centrifuged another 30 seconds at full speed to collect and remove residual ethanol. The samples were air dried, resuspended in 40 µL RNase-free water. 20 µL was frozen at −80ºC for longer term storage, and 20 µL was used for quantification and stored at 20ºC for subsequent cDNA synthesis. RNA concentrations were measured with a Qubit XR assay kit (Invitrogen, Q33224). cDNA was synthesized with the iScript Reverse Transcription Supermix for RT-qPCR (Bio-Rad, 1708841), following the manufacturer’s protocol. Real-time PCR was conducted with the iTaq Universal SYBR Green Supermix kit (Bio-Rad, 1725121) using the manufacturer’s protocol and reaction parameters for the QuantStudio3 (Applied Biosystems), and an annealing temperature of 60ºC for all primer sets. All primers were designed to anneal to neighboring exons to prevent amplification from genomic DNA (**S1 Table**). In addition to 24 samples, each plate contained a standard curve, a calibrator sample, and two negative controls (a cDNA reaction without reverse transcriptase and water instead of sample). Each biological sample was run in technical triplicate.

Expression data (Cp values) were exported from the QuantStudio3 software and analyzed in R. For each triplicate, outliers were excluded if the crossing point values (Cp) differed by more than 0.5 cycles and the standard deviation was over 0.2. After exclusion of outliers, the average Cp value was calculated per sample and used for further analysis. The standard curves were used to calculate relative concentrations for each sample and gene based on the Cp. To calculate a normalized expression (ΔC_T_) the concentration of the target gene was divided by the concentration of the endogenous control *rpl-26* [31]. To calculate the relative normalized expression (ΔΔC_T_) between different plates of the same gene, the normalized expression of the samples was divided by the normalized expression of the calibrator. The effect of treatment (PZQ or control) and strain (N2 and JU775) on expression of *cct-8, hsp-16*.*2*, and *hsp-70* was tested with a two-way ANOVA, including the interaction between treatment and strain and the effect of plate (R package *stats*, version 4.1.2). Before analysis, outliers were excluded by removing any values above or below two standard deviations from the mean. For *cct-8*, a plate-effect was found, and both plates were subsequently analyzed independently.

### 2.6 qPCR experiment to determine *hsp* induction after RNAi treatment

A population of the N2 strain was grown as previously described and grown on bacterial plates containing *E. coli* expressing: L4440 empty vector, *act-1* RNAi, *cct-8* RNAi, or *act-1* and *cct-8* RNAi. The populations were grown for 48 hours and then placed in 2% DMSO or 4 mM praziquantel (2% DMSO) for eight hours. RNA was then extracted using a standard Trizol and chloroform extraction [32]. Purified RNA was treated with DNAse I, pelleted, and resuspended in water. RNA was synthesized into cDNA using first-strand synthesis (Bio-Rad iScript Advanced cDNA Synthesis Kit for RT-qPCR, 1725037). qPCR was then performed as previously described [33] using custom primers (**S1 Table**). Amplification of the 18S region was used to ensure the measurements were in the linear range and amplification of *hsp-16*.*2* transcripts were used to measure heat-shock induction.

### 2.7 Generation of genome-edited strains

Genome-edited allele replacement strains were generated with a *dpy-10* co-CRISPR strategy [34]. ALT-R and tracrRNA were purchased from IDT (Skokie, IL) (**S2 Table**). The tracrRNA (IDT, 1072532) was injected at a final concentration of 13.6 µM. The crRNA for *dpy-10* was injected at 4 µM, and the crRNA for *cct-8* was injected at 9.6 µM. The repair single-stranded oligonucleotide repair templates were injected at 1.34 µM for *dpy-10* and 4 µM for *cct-8*. Purified Cas9 protein (IDT, 1074182) was injected at final concentration 23 µM. Injection mixtures were generated by combining the tracrRNA and crRNAs and incubating them at 95ºC for five minutes and 10ºC for 10 minutes. Then, Cas9 was added and an additional five-minute room temperature incubation was performed. Following these incubations, repair templates and nuclease-free water were added to the injection mixture. This mixture was then loaded into a pulled microinjection needle (World Precision Instruments, 1B100F-4). Injected animals were singled onto 6 cm NGMA plates and allowed to produce offspring. These offspring were screened for Rol or Dpy phenotypes. Individuals showing the desired phenotypes were singled to new 6 cm NGMA plates and allowed to produce offspring. Then, the *cct-8* region (oECA1155 and oECA1157) of the F_1_ Rol or Dpy parents were amplified using PCR. The PCR product was digested with the *Nla*IV restriction enzyme and differential restriction patterns were identified. Parents with the desired restriction pattern were then Sanger sequenced to verify homozygous edits, and the *dpy-10* mutation was crossed out of the strain. Custom primers, guides, and repair templates were used (**S2 Table**).

### 2.8 Data availability

The data and code to replicate analysis and figure generation are available at https://github.com/AndersenLab/Praziquantel.

## 3. RESULTS AND DISCUSSION

### 3.1 The racemic mixture and both enantiomers of PZQ affect *C. elegans* development

To test if PZQ has an effect on the nematode *C. elegans*, we assayed nematode development in response to racemate PZQ. Development was measured for thousands of animals using a previously developed high-throughput assay (HTA) (see 2.2) [15,24–26,35–40]. In summary, three L4 larvae were sorted into each well of a 96-well plate in the presence of DMSO or PZQ dissolved in DMSO. In the next 96 hours, these larvae grew and reproduced, and their offspring developed. After 96 hours, animal length (time of flight, TOF) and optical density (extinction, EXT) were measured for every animal of the population in each well as a proxy for development [41]. Animals grow longer and more dense over time, and toxic drug treatments slow this development. Therefore, if PZQ is detrimental to *C. elegans*, animals are expected to be shorter and less optically dense after 96 hours. Here, we report the median optical density normalized for animal length as the phenotype (median.norm.EXT). At doses of 1 mM and higher, PZQ inhibits development of four genetically distinct strains (**S1 Fig**.). The Hawaiian strain CB4856 is sensitive to PZQ treatment compared to the laboratory strain N2, whereas the Portuguese strain JU775 is less affected than the N2 strain. Although PZQ is administered to both humans and animals as a racemic mixture, only the left-handed PZQ ((*S)*-PZQ) is effective at fighting schistosome infections *in vivo* and *in vitro* [reviewed in 42]. In a subsequent assay, the N2 and JU775 strains were subjected to racemate PZQ as well as to both enantiomers. The enantiomers are equally as potent as the racemate and inhibited development of both strains up to a dose of 1 mM, after which the enantiomers were slightly less potent than the racemate, but no difference was found between the two enantiomers (**Fig 1, Tables S3 and S4**). The schistosome drug target *Sm*.TRPM_PZQ_ has no known orthologs in *C. elegans* [43,44], which in combination with enantiomer activity could imply a different mode of action in nematodes, suggesting a reason why both enantiomers might be active. Further studies on the mode of action and drug target of PZQ in natural populations of both species will determine the relevance of *C. elegans* as a model for schistosome responses to PZQ. Regardless, the reported effectiveness of PZQ in inhibiting development of *C. elegans* suggests optimization of PZQ could be explored for anthelmintic treatment of nematodes.

**Fig 1:**
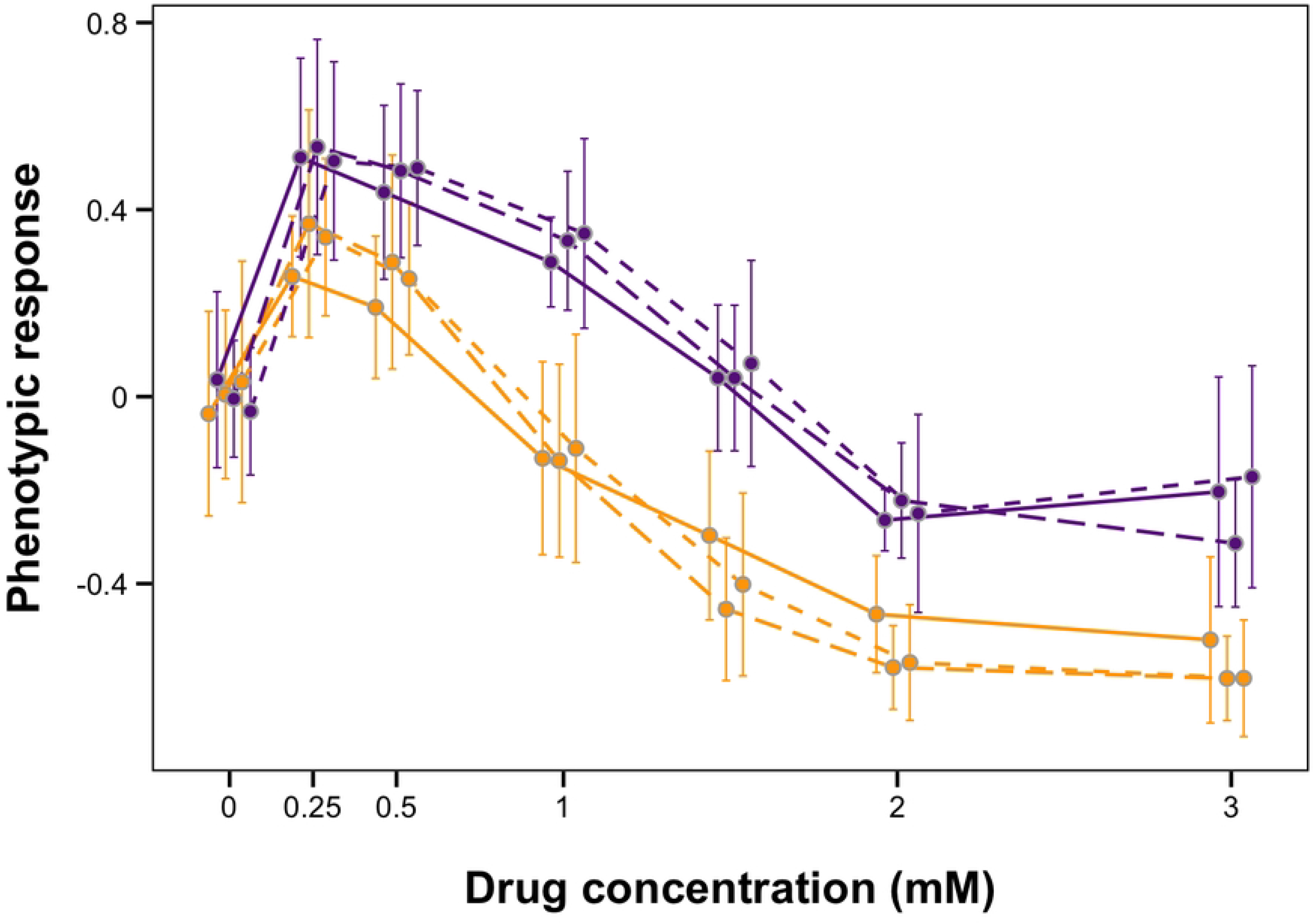
Praziquantel affects C. elegans development. Dose response for the N2 (orange) and JU775 (purple) strains in praziquantel (PZQ), (S)-PZQ, and (R)-PZQ. The phenotypic response represents median optical density normalized for animal length. Linetype indicates type of drug: solid = PZQ racemate, long dash = (S)-PZQ, and short dash = (R)-PZQ.

### 3.2 Genome-wide association mapping of 74 wild isolates implicates a region on chromosome IV in the response to PZQ

The observed variation in responses to PZQ across four wild strains (**S1 Fig**) suggested that genetic variation of these strains could underlie the differential responses. To further study the effect of genetic variation on PZQ susceptibility, development of 74 wild *C. elegans* strains was quantified in 1 mM of PZQ in 1% DMSO or in control conditions (1% DMSO) with the same HTA as described for the dose responses. Because the PZQ racemate and its enantiomers had similar effects at 1mM in the dose response assay, the racemate was used in this experiment. We performed a genome-wide association (GWA) mapping and identified a quantitative trait locus (QTL) on the left arm of chromosome IV (position 845,848 to 1,313,281) (**Fig 2A**). In this set of 74 strains, the N2 strain was more susceptible than the JU775 strain, as was observed in the initial dose response (**Fig 2B, S1 Fig**.). Fine mapping of this region shows 58 genes are contained in the QTL, of which 15 genes have a −log_10_(*p*) of over 10, and five genes a −log_10_(*p*) greater than 13.5 (**S5 Table, Fig 2C**). Two of these genes, *cct-8* and *znf-782*, have missense variants in the JU775 strain, respectively G226V (*cct-8*), and Q617H and H563D (*znf-782*). The missense variant in *cct-8* is likely to affect protein function. This G226V variant was found in many wild *C. elegans* strains, including JU775, suggesting that the valine at position 226 might reduce PZQ sensitivity. *cct-8* encodes a subunit of the TRiC complex, which has essential roles in the folding of a large number of proteins, including actin [45].

**Fig 2:**
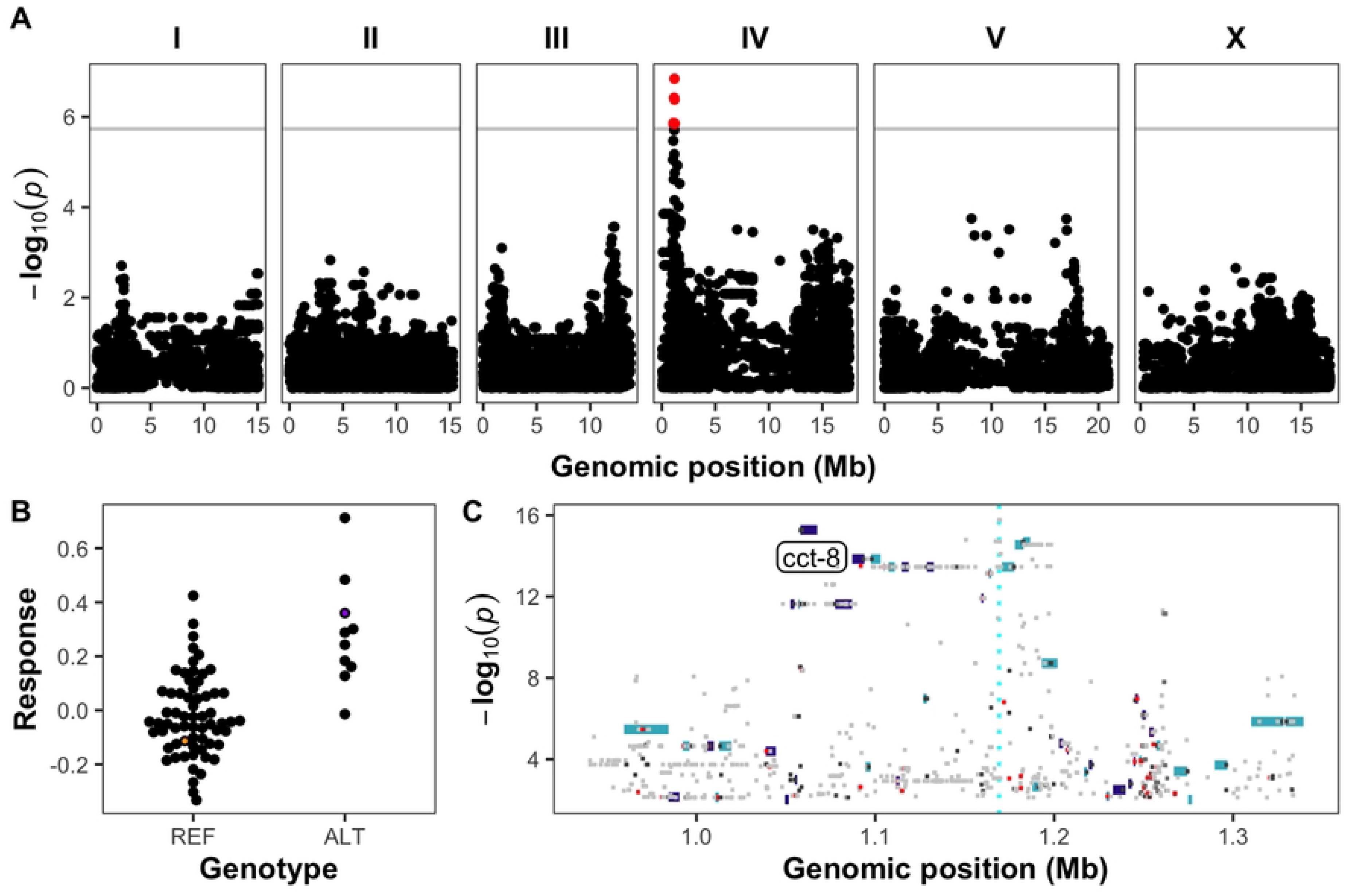
A) Genome-wide association mapping results for median normalized optical density (median.norm.EXT) across 74 wild strains. The genomic position (x-axis) is plotted against the −log_10_(p) value (y-axis) for each SNV. Legend: dots: SNV, red dot: SNV passes the genome-wide Bonferroni significance threshold designated by the gray line. **B)** At the SNV with the highest −log_10_(p) value (position 1,169,239), genotypes are split based on the presence of the reference allele (REF) or the alternative allele (ALT). The y-axis shows the normalized optical density (median.norm.EXT) with each point representing the average median.norm.EXT response of a single wild strain. The N2 strain is shown in orange, and the JU775 strain is shown in purple. **C)** Fine mapping of variants in the QTL on chromosome IV. The genomic position is shown on the x-axis, and the −log_10_(p) value for variants is shown on the y-axis. Genes within the region are shown as rectangles and colored by the strand (turquoise = + strand, purple = - strand). Variants (amino acid changes) with a predicted moderate impact on gene function are shown in orange, and variants with a predicted high impact on gene function (frame-shift, stop-gain, start-loss) are shown in red. The gene cct-8 is labeled on the plot directly left of the corresponding rectangle for the gene.

### 3.3 Expression analysis of *cct-8* further implicates it in the *C. elegans* response to PZQ

The TRiC complex is essential for the proper folding of actin and many other cellular proteins. Disruption of the TRiC complex leads to actin misfolding and activation of a cellular pathway known as the heat-shock response [46]. To investigate if the *C. elegans* response to PZQ involved the disruption of the TRiC complex, we first measured the expression of *cct-8, hsp-16*.*2*, and *hsp-70* in the N2 and JU775 strains in control conditions and after PZQ treatment (1 mM PZQ in 1% DMSO), with the expression of *rpl-26* serving as a control. A significant effect of strain but not condition was detected for *cct-8* expression (**Fig 3A, S6 Table**). Expression was higher in the less susceptible strain JU775 both with and without treatment. Expression of *hsp-16*.*2* did not differ between strains or conditions, and *hsp-70* was expressed differentially between the strains but again not affected by treatment (**Fig 3A, S6 Table**). These data imply that the JU775 *cct-8* allele (G226V) requires higher expression in order to elicit sufficient function of the TRiC complex. This higher expression of *cct-8* in turn might lead to internal stress and an upregulation of *hsp-70*, although PZQ treatment itself does not promote further upregulation of this stress response.

**Fig 3:**
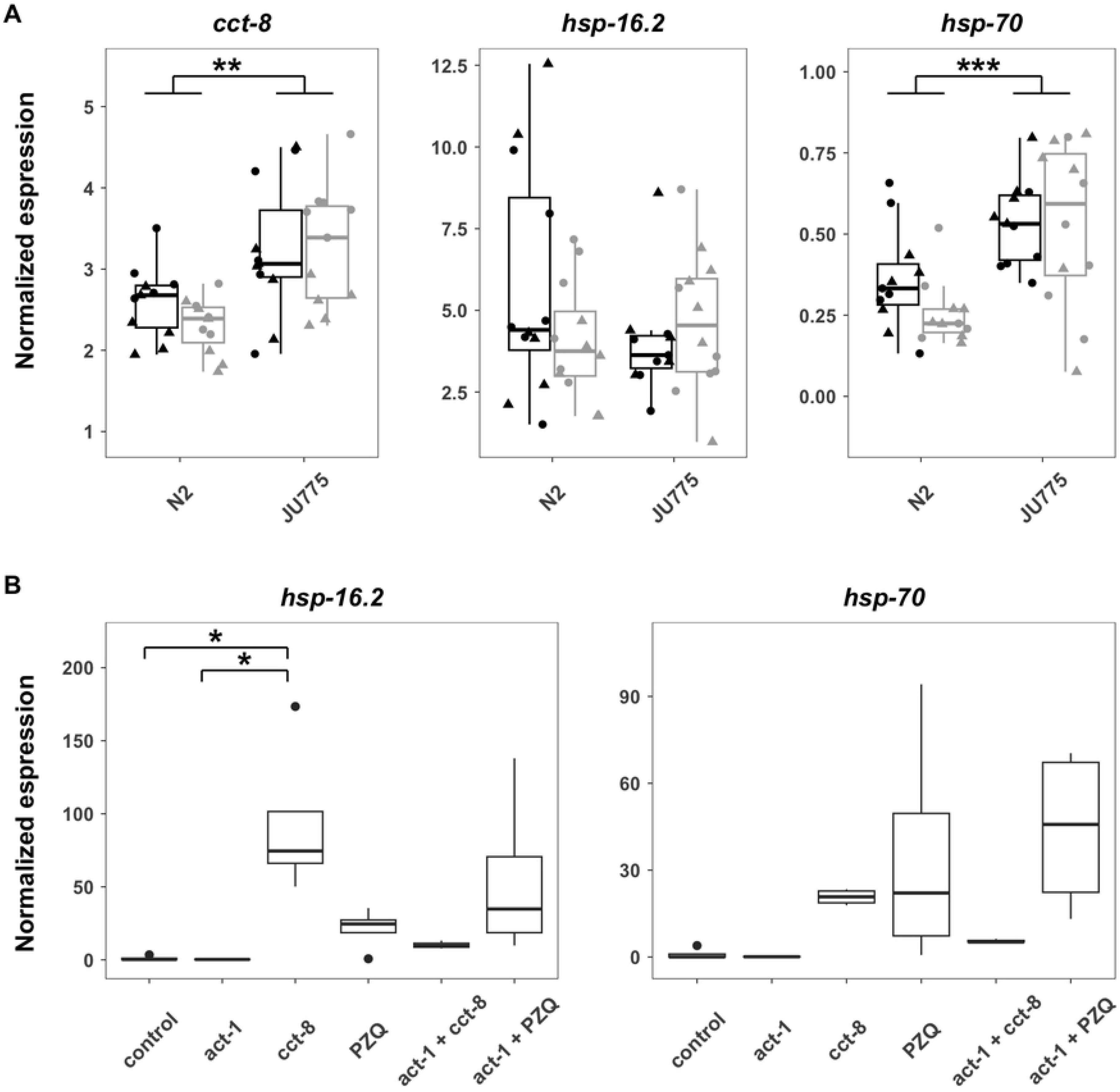
A) Expression of cct-8, hsp-16.2, and hsp-70 in control conditions (black) and PZQ (gray) in the strains N2 and JU775. **B)** Expression of hsp-16.2 and hsp-70 after treatment with RNAi or praziquantel (PZQ) in the strain N2. *, **, and *** indicate p-values of less than 0.05, 0.01, and 0.001, respectively. The biological samples were spread over two 96-well plates per gene (triangles and circles).

Next, we tested if acute PZQ exposure at a higher dose (4 mM PZQ in 2% DMSO) for eight hours affects *hsp* expression in the N2 strain [47]. We observed that individuals showed a significant increase in expression of *hsp-16*.*2* after *cct-8*-RNAi treatment and a non-significant increase after PZQ treatment (**Fig 3B**). Furthermore, the knockdown of *act-1* to reduce the level of misfolded proteins in combination with RNAi of *cct-8* or PZQ treatment had a less severe heat-shock response than *cct-8* knockdown or PZQ treatment alone (**Fig 3B**). This trend offered some support to our hypothesis that PZQ might alter the function of TRiC and that *cct-8* might play a role in PZQ sensitivity.

Finally, we looked at expression of genes in the QTL on chromosome IV to see if mediation might affect the response to PZQ. Mediation analysis tests if expression of the genes in a QTL is mediated by genomic variation either close to (local eQTL) or farther away from (distant eQTL) the focal gene. None of the genes in this region, including *cct-8*, showed significant mediation (**S5 Table**). These data suggest that the regulation of gene expression by genetic variation elsewhere in the genome did not contribute to the differences in PZQ responses across the 74 strains studied here.

### 3.4 Variation at *cct-8* codon 226 does not explain differential PZQ responses between the N2 and JU775 strains

To test the hypothesis that the G226V allele is involved in reduced sensitivity to PZQ, we used CRISPR-Cas9 genome editing to introduce the V226 allele into the N2 genetic background and the G226 allele into the JU775 genetic background (**Fig 4A,B**). We then measured PZQ responses for both parental strains and both genome-edited strains using the high-throughput assay. We found that this variant does not underlie the phenotypic differences between the N2 and JU775 strains (**Fig 4C**). We looked for gene duplications or deletions in the JU775 strain relative to the N2 strain as an alternate source for variation in the TRiC complex, but found no evidence of this copy-number change. Taken together with the various types of expression data, *cct-8* is a candidate for PZQ mode of action and mechanisms of resistance in *C. elegans*, but further studies are needed to reach a more definitive conclusion.

**Fig 4:**
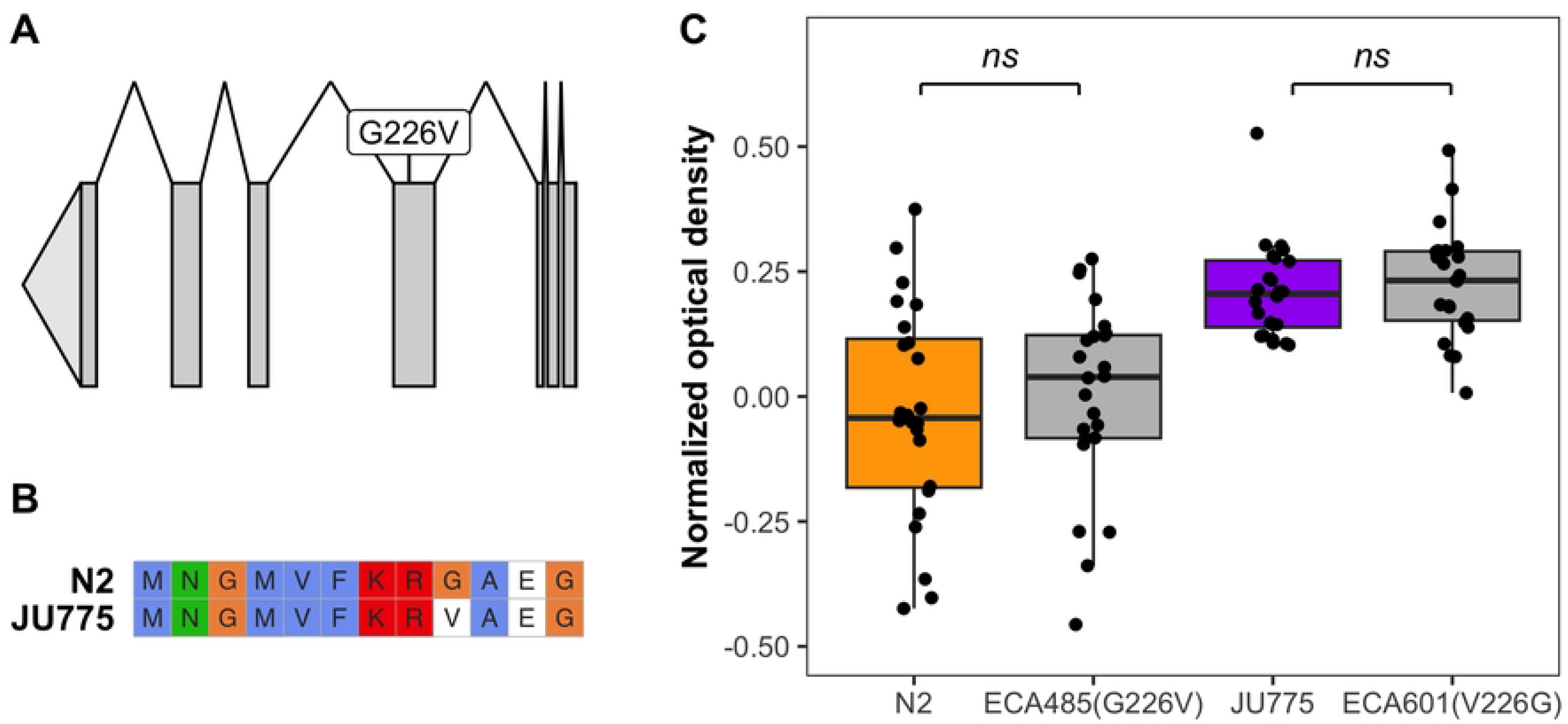
A) The gene model for the longest isoform of cct-8 is shown. Exons are shown in dark gray and introns as lines connecting the exons. The variant G226V is shown on the exon where it is located. **B)** The amino acid sequences of the N2 and JU775 versions of CCT-8 are shown aligned around the region containing the G226V variant. Figure made with ggmsa [48] **C)** The x-axis shows the strains tested: N2, ECA485 (N2 background with the JU775 allele in cct-8), JU775, and ECA601 (JU775 background with N2 allele in cct-8). Relative normalized optical density (median.norm.EXT) in 1 mM praziquantel is shown on the y-axis. Each point represents the median.norm.EXT for a population of animals in the presence of praziquantel normalized for their response in 1% DMSO. ns = not significant.

### *3*.*5 C. elegans* as a potential model for PZQ responses in schistosomes

The CCT/TRiC pathway identified in this study as potentially involved in PZQ responses is conserved across eukaryotes [49]. Further study of this pathway in *C. elegans* might yield interesting hypotheses for schistosomes studies. With the lack of orthologs of the schistosome drug target *Sm*.TRPM_PZQ_ in *C. elegans* [43,44], no clear connection can be made to test this mechanism using *C. elegans* as a model for PZQ responses at the present time. However, as noted by Cotton and Doyle [6], the *Sm*.TRPM_PZQ_ was found in one genetic background [11,50]. In this study, we started with 74 wild strains, which by no means captures the variation present in the global *C. elegans* population [51,52]. It is therefore conceivable that other molecular targets or modes of action exist in either species and that orthologous genes or pathways are present [7]. Furthermore, the trait used to determine sensitivity can affect strain responses [40,53] and association mapping results. Maybe development is not the ideal trait to study in *C. elegans*, because PZQ efficacy is age-dependent and less active against juvenile schistosomes [54,55] and studying lethality of adult nematodes might be a better comparison. Regardless of its use for schistosome PZQ studies, a better understanding of PZQ action in *C. elegans* could help optimize PZQ as an anthelmintic for nematodes, capitalizing on its arsenal of high-quality wild strain genomes and well developed protocols for genome-editing and genetic crosses, amongst other techniques.

## 4. CONCLUSIONS

In this work, we describe the effect of PZQ on development of *C. elegans*, and the potential mechanisms underlying resistance to PZQ in this nematode model. We explore genome-wide variation to identify a genomic region on chromosome IV that is correlated with resistance, but both genome engineering and gene expression of the candidate gene *cct-8* do not provide a conclusive hypothesis for the mode of action or resistance mechanism of PZQ. We propose that TRiC function in the resistant strain JU775 is negatively affected, potentially caused by a mutation in *cct-8*. This decreased function is compensated by increased expression of *cct-8*, which causes higher baseline levels of internal stress as shown by increased *hsp-70* expression. The increased expression of *hsp-70* in turn makes the JU775 strain more resilient to the stress caused by subsequent PZQ treatment, making it more resistant.

## ACKNOWLEDGMENTS

We would like to thank Savannah Brennan for her support with this project and members of the Andersen laboratory for their feedback throughout the project. We would also like to thank Kristina Kraml and Lotus Separations, LLC for providing the PZQ enantiomers, as well asWormbase, the *Caenorhabditis* Genetics Center (P40 OD010440), and the *Caenorhabditis* Natural Diversity Resource (NSF Capacity 2224885).

## SUPPORTING INFORMATION CAPTIONS

**S1 Table. Primer sequences for qPCR**

**S2 Table**. Oligonucleotide sequences for generation of ECA485 *cct-8*(*ean8*) in a N2 background and ECA601 *cct-8*(*ean39*) in a JU775 background.

**S3 Table**. Two-way ANOVA results for dose response assay of racemate praziquantel (PZQ), (*S*)-PZQ, and (*R*)-PZQ.

**S4 Table**. *post hoc* Tukey HSD Test for the effect of drug types (racemate praziquantel (PZQ), (*S*)-PZQ, and (*R*)-PZQ) on development at doses 1500, 2000, and 3000 mM.

**S5 Table**. Genes found in the fine mapped region on chromosome IV, position 939,925 to 1,334,212 in order of descending significance.

**S6 Table**. Two-way-ANOVA results for *cct-8, hsp-16*.*2*, and *hsp-70* expression in the strains N2 and JU775 in control (1% DMSO) and PZQ conditions (1mM PZQ in 1% DMSO).

**S1 Fig**. The effect of increasing concentrations of Praziquantel (PZQ) on *C. elegans* development.

